# DDS-E-Sim: A Transformer-based Probabilistic Generative Framework for Simulating Error-Prone DNA Sequences for DNA Data Storage

**DOI:** 10.1101/2025.02.14.637785

**Authors:** Mst. Fahmida Sultana Naznin, Swarup Sidhartho Mondol, Adnan Ibney Faruq, Debashmita Saha, Ahmed Mahir Sultan Rumi, A. B. M. Alim Al Islam

## Abstract

DNA has emerged as a promising medium for long-lasting data stoage due to its high information density and long-term stability. However, DNA storage is a complex process where each stage introduces noise and errors. Since running DNA data storage experiments in vitro is still expensive and time-consuming, a simulation model is quite necessary that can mimic the error patterns in the real data and simulate the experiments. Existing tools often rely on fixed error rates or are specific to certain technologies. We propose DDS-E-Sim, a transformer-based probabilistic generative framework that simulates errors in a DNA data storage channel, regardless of the process or technology. DDS-E-Sim successfully captures the error distribution of DNA storage pipelines and learns to stochastically generate erroneous DNA reads. Given oligos (DNA sequences to write), it outputs erroneous reads resembling real pipelines capturing both random and biased errors, such as k-mer and transition errors. Evaluations on two distinct technology-specific datasets show high fidelity and universality: DDS-E-SIM exhibit a total error rate deviation of only 0.1% and 0.7% respectively on the datasets processed with Illumina MiSeq and Oxford Nanopore. Additionally, our simulator generates 100,743 unique oligos from 35,329 sequences, with coverage 5 (each sequence read five times) in the test datasets, demonstrating its ability to simulate biased errors and stochastic properties simultaneously.

## 1 Introduction

In the era of data expansion, the world produces 10^18^ bytes daily [26], demanding durable storage (*>*50 years) [41]. DNA offers high information density and stability with minimal maintenance power [11]. DNA Data Storage (DDS) comprises multiple stages - synthesis, storage, sampling, and sequencing, each introducing potential insertion, deletion, and substitution errors[6, 39]. DDS errors are both synchronous and asynchronous with complex statistics [20]. Despite reduced synthesis and sequencing costs [28, 10, 5], large-scale DDS remains costly. Simulation provides a low-cost, probabilistic approach for channel verification, synthesis analysis, and ECC evaluation [14, 22, 23, 4] by modelling synthesis imperfections, molecular decay, PCR biases, and sequencing noise [2, 9].

Numerous prior DDS studies characterize single-stage error simulation separately [38, 21, 29, 27], but fail to capture end-to-end error evolution. Existing simulators such as ART [16], Flux [12], and pIRS [15] focus on specific sequencing stages. Nanopore-focused tools (DeepSimulator [24, 25], NanosigSim [7], NanoSim [30]) remain technology-specific. DeSP [39] is a flexible model which can adapt to diverse experiment conditions but it uses fixed user-defined parameters. WGAN [20] is a recently developed universal simulator that is not process dependent but it struggles with longer Nanopore sequences because of its GAN architecture. Moreover, extensive hyperparameter tuning is required for stable GAN training.

Conventional DNA data storage error modelling involves numerous challenges. Firstly, Errors in DNA data storage display both systematic biases and stochastic behaviour [37, 20]. Most of the existing simulators rely on predefined, hard-coded probabilities [31], which overlook stochastic variability. Conversely, some of the models may achieve that but fail to account for systematic biases [32, 3]. We address the problem with a Transformer-based generative model within a Beta-VAE framework that additionally incorporates Gaussian noise for stochasticity and retains dominant biased patterns. Secondly, error behaviours differ by technology[15, 24]. For instance, Illumina uses fluorescence for short, precise reads, while Nanopore detects electrical signals for long, relatively more error-prone reads. Most DDS simulators remain technology-specific. Researchers need to switch simulators at different stages or datasets to simulate end-to-end errors, as error profiles vary with technology and even with system configurations, which creates a major bottleneck. This highlights the need for a universal simulator that integrates multiple stages and technologies. A unified framework would streamline diverse DDS methods and facilitate system design. Our framework is data-centric rather than process-centric, and its validation on diverse technological datasets ensures robust cross-process generalization. In summary, our main contributions are as follows:

- We introduce a novel transformer-based probabilistic generative model within a Beta-VAE framework for error simulation in DDS. We additionally incorporate a Gaussian noise–based perturbation to further enhance stochasticity, which successfully introduces randomness and aligns the generated samples with the statistical distribution of the datasets. Our simulator is universal and simulates erroneous DNA reads regardless of technology.
- We successfully demonstrate the robustness of our data-centric universal modelling by simulating error profiles from two distinct datasets across different technologies. We conduct extensive experiments to quantify insertion, deletion, and transition error rates, as well as k-mer error patterns. Our simulator generates DNA reads that closely match real error profiles observed in the test datasets and outperforms existing tools.

## 2 Methods

We use a Transformer within a Beta-VAE framework to learn disentangled error representation in DNA sequences. An overview of the architecture is shown in Figure 1.

**Figure 1.**
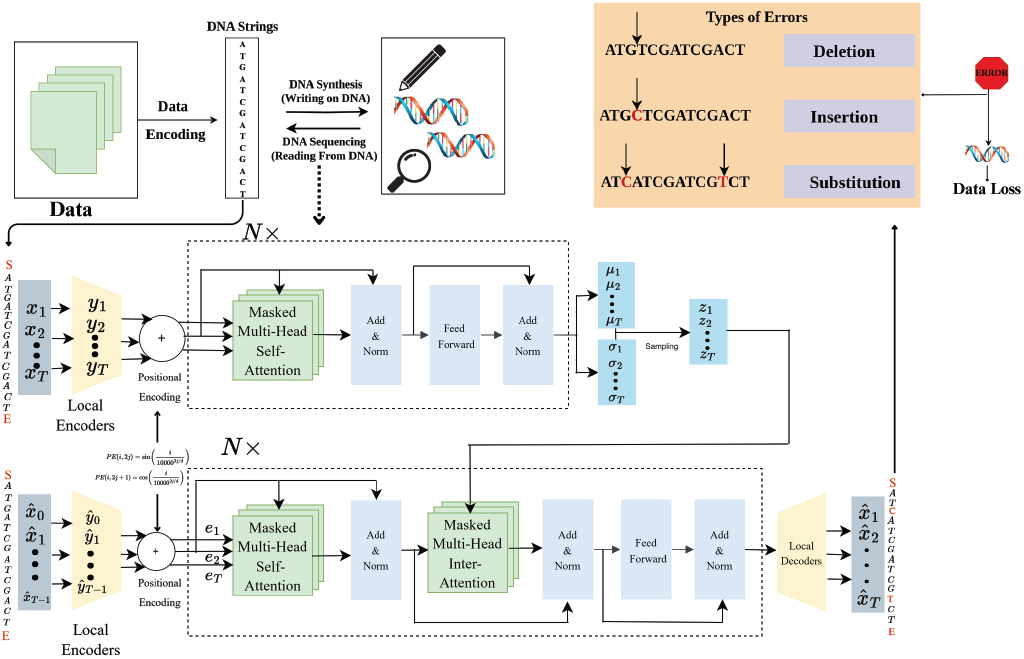
An overview of DDS-E-Sim architecture and DNA data storage pipeline.

### 2.1 Problem Formulation

The goal of the model is to replicate the sequential processes of the DDS, from the input oligo sequence to the output read sequence. Let *X* = {*A, C, G, T* } represent the set of nucleotide bases, and *X*_*e*_ = {*A, C, G, T, S, E, P*} represent the set of extended symbols, where ‘S’ is the start marker, ‘E’ is the end marker, and ‘P’ is padding. The input to the simulator is a nucleotide sequence, denoted *x* = *x*_1_*x*_2_ … *x*_*N*_, with *x*_*i*_ ∈ *X*_*e*_ for 1 ≤ *i* ≤ *N*. The output of the simulator is an error-prone DNA sequence 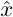, referred to as the read, where 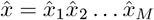, with 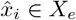 for 1 ≤ *i* ≤ *M*. Initially, sequence length variation was addressed using padding but we later adopted a sliding-window strategy, which proved to be a slightly more effective choice.

### 2.2. Proposed Stochastic Generative Learning

#### Inducing Stochasticity with Gaussian Noise in DNA Sequences

In DDS, error-prone sequences must reflect stochasticity across coverage levels. We represent characters through one-hot encoding and manage long sequences by first applying padding, then switching to a sliding-window strategy sliding over the sequence with window and step size. We add Gaussian noise to model synthesis and sequencing variability as unbiased synthesis and sequencing processes naturally exhibit Gaussian-like behaviour [8, 37]. Noisy sequences are generated as *y* = *x* + *N* (*µ, σ*^2^), where *x* is the original sequence, *y* the perturbed sequence, *µ* is the mean (typically 0), and *σ*^2^ is the variance, which controls the intensity of the noise. Through a random search experiment, we determine an optimal noise factor.

#### Transformer Architecture for Capturing Error Patterns

Our model integrates a Transformer [34] in a Beta-VAE framework [13] to learn error patterns from DNA sequences. We incorporate masked and causal attention to preserve sequential dependencies [19]. Let *x*_1..*T*_ be a DNA sequence of length *T*, where *x*_*i*_ denotes the *i*-th nucleotide. After noise addition, each nucleotide passes through a local encoder 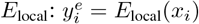 yielding nucleotide-level representations 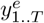. The Transformer encoder computes the latent parameters:

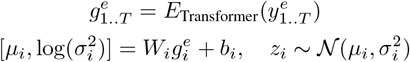

The latent codes are decoded sequentially using the Transformer decoder *D*_Transformer_ and local decoder *D*_local_:

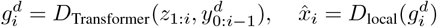

Here 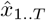 is the generated error-prone sequence, and 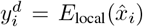. A special embedding 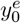 denotes the start of sequence, analogous to the SOS token in Natural Language Processing.

#### Causal Masking for Sequential Cohesion Control (CM)

In DDS, nucleotide sequences must follow a strict order to ensure correct encoding and retrieval. Similar to biological transcription and sequencing, the model must not access future positions, as order disruption leads to retrieval errors [36, 35]. To emulate this, we apply causal masking across encoder and decoder layers, enforcing directionality so that the *i*-th base attends only to positions *j* ≤ *i*. The masked attention is defined as

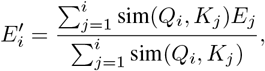

where 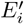 is the updated representation of base *i, Q*_*i*_ and *K*_*j*_ are query and key vectors, *E*_*j*_ is the base representation, and sim(*Q*_*i*_, *K*_*j*_) denotes their similarity score.

#### Balancing Error Bias Modelling and Stochasticity with Beta-VAE

DNA sequences are inherently complex because they contain hierarchical and structured dependencies [17]. For example, certain genes are organized into clusters on the DNA strand, such as the Hox gene cluster, which is essential for the coordinated expression of genes during development [17]. Additionally, regulatory elements like enhancers, silencers, and promoters interact in a hierarchical manner to control gene expression [40]. Thus, errors in DNA sequences inherently carry biases and exhibit dominance in certain regions or motifs [1]. Repetitive sequences or regions with high GC content causes sequencing errors. The model needs to capture error patterns inherent in DNA sequences, such as k-mer biases, positional skewness, and dominance of specific transitions. At the same time, it should introduce stochasticity to ensure diverse and realistic error representations. We use Beta-VAE with transformer architecture to achieve that. The objective function can be expressed as:

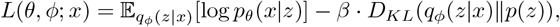

Here, 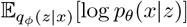 is the expected log-likelihood of the data under the approximate posterior distribution *q*_*ϕ*_(*z* |*x*), *D*_*KL*_(*q*_*ϕ*_(*z* |*x*) ∥ *p*(*z*)) is the Kullback-Leibler divergence between the approximate posterior *q*_*ϕ*_(*z* |*x*) and the prior *p*(*z*). Here, the value of *β* is of utmost importance. We observe that high value *β* over-regularizes the latent space, leading to under-representation of motif-dependent DDS error patterns. A moderate setting of *β* = 0.5 preserved the essential error dependencies while maintaining sufficient regularization to prevent overfitting.

## 3 Experiments and Results

We conduct extensive experiments on two datasets from different sequencing platforms and compare our model against established baselines. We analyze insertion, substitution, and deletion error rates, as well as k-mer error patterns and positional error distributions, to comprehensively assess how well the models capture the inherent error characteristics of DNA data storage pipelines.

### 3.1 Dataset Description

We evaluate two technology-specific datasets to assess universality. The first dataset [18] has 18,000 oligos (length 152, GC 45–55%, homopolymer ≤ 3) sequenced via Illumina (15,126,429 reads; train: 12,108,573, test: 2,999,656). The second dataset [33] has 10,000 oligos (length 110) sequenced via ONT MinION (269,709 reads; train: 242,738, test: 26,971). We represent datasets D-I (Illumina) and D-N (Nanopore) respectively.

### 3.2 Performance Evaluation

We conduct an ablation study with three variants: (1) **Transformer+Beta-VAE:** our proposed model, **Transformer**: excluding VAE from the our architecture, and (3) **VAE:** a Multi-Attention-LSTM with three LSTMs and a VAE (using LSTM instead of transformer in our architecture). In addition, we compare against **WGAN** [20], the only universal deep learning simulator we are aware of. By contrast, simulators such as DeepSimulator [25] and Illumina [16] are technology-specific and cannot be applied across datasets from different sequencing platforms.

#### Uneven Base-Level Error Statistics

We evaluate single-base errors against original data and baselines in Table 1. For D-I, substitution is the dominant form of error. DDS-E-Sim shows the smallest deviations from original reads and outperforms others by a good margin. Dominant transitions are well captured in our proposed model as shown in Table 2a. DDS-E-Sim also achieves minimal deviations For D-N and accurately models base transitions as shown in Table 2b. DDS-E-Sim achieves an impressive total error deviation of only 0.1% and 0.7% in the two datasets.

**Table 1.**
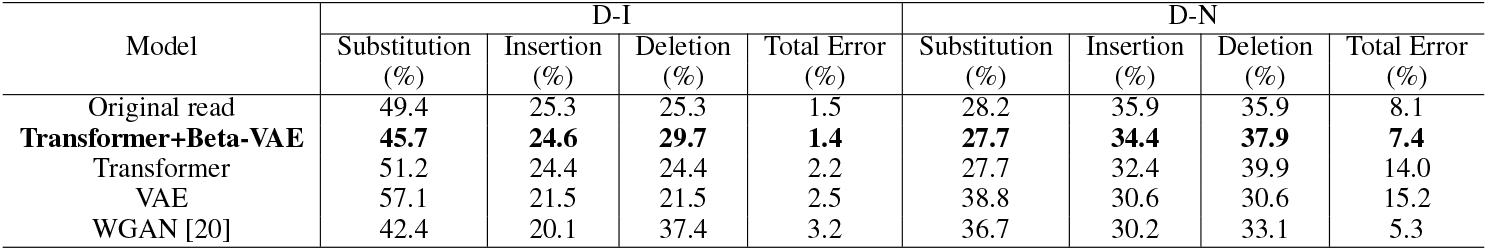
Error rates of original and model-generated DNA reads across datasets D-I and D-N.

**Table 2.** Transition error rates across datasets D-I and D-N. Each row shows relative error rates of other bases for the base being substituted from. For each transition, the error rate of the best performing model (least deviation from the original read) is shown in bold.

| (a) Dataset D-I |  |  |  |  |  |  |  |  |  |  |  |  |
| --- | --- | --- | --- | --- | --- | --- | --- | --- | --- | --- | --- | --- |
|  | Original read |  |  |  | DDS-E-Sim |  |  |  | WGAN |  |  |  |
|  | A | C | G | T | A | C | G | T | A | C | G | T |
| A | – | 0.47 | 0.36 | 0.17 | – | **0.42** | **0.39** | **0.19** | – | 0.30 | 0.30 | 0.39 |
| C | 0.35 | – | 0.31 | 0.34 | **0.68** | – | **0.15** | **0.17** | 0.23 | – | 0.06 | 0.71 |
| G | 0.38 | 0.10 | – | 0.53 | 0.54 | **0.12** | – | 0.34 | **0.36** | 0.07 | – | **0.57** |
| T | 0.33 | 0.18 | 0.49 | – | **0.36** | **0.18** | **0.45** | – | 0.22 | 0.59 | 0.20 | – |

| (b) Dataset D-N |  |  |  |  |  |  |  |  |  |  |  |  |
| --- | --- | --- | --- | --- | --- | --- | --- | --- | --- | --- | --- | --- |
|  | Original read |  |  |  | DDS-E-Sim |  |  |  | WGAN |  |  |  |
|  | A | C | G | T | A | C | G | T | A | C | G | T |
| A | – | 0.22 | 0.58 | 0.19 | – | **0.22** | **0.57** | **0.21** | – | 0.30 | 0.29 | 0.41 |
| C | 0.21 | – | 0.17 | 0.62 | **0.21** | – | **0.17** | **0.62** | 0.16 | – | 0.16 | 0.68 |
| G | 0.61 | 0.18 | – | 0.22 | **0.59** | **0.17** | – | **0.24** | 0.23 | 0.22 | – | 0.55 |
| T | 0.19 | 0.61 | 0.20 | – | **0.20** | **0.59** | **0.21** | – | 0.22 | 0.53 | 0.25 | – |

#### Uneven k-mer Error Patterns

We analyze 2-mer and 3-mer error rates to assess motif-specific biases (Table 3). In dataset D-I, our model reproduces the expected trends: 3-mer errors exceed 2-mer errors, with AAA being the most error-prone motif and TT the least. In dataset D-N, k-mer analysis shows CCC as the most error-prone motif and AA as the least, with 3-mer errors again higher than 2-mers, thereby preserving higher-order error patterns. Overall, the model effectively captures both motif- and length-dependent error characteristics.

**Table 3.**
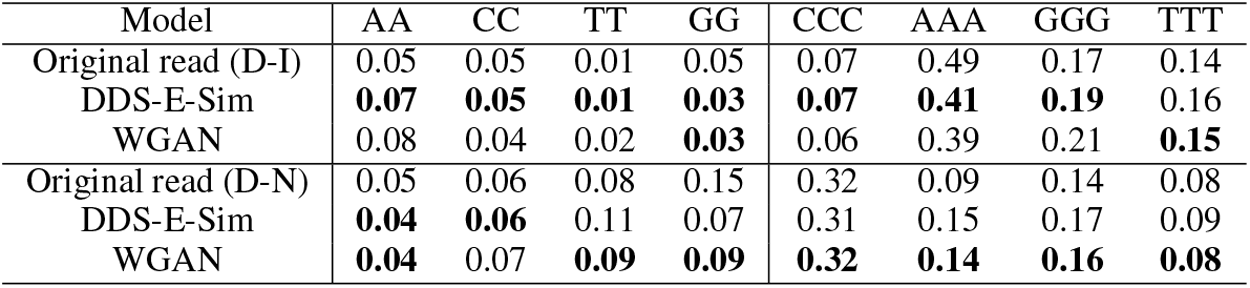
K-mer error rates of different models across datasets D-I and D-N. For each k-mer, the error rate of the best performing model (least deviation from the original read) is shown in bold.

| Model | AA | CC | TT | GG | CCC | AAA | GGG | TTT |
| --- | --- | --- | --- | --- | --- | --- | --- | --- |
| Original read (D-I) | 0.05 | 0.05 | 0.01 | 0.05 | 0.07 | 0.49 | 0.17 | 0.14 |
| DDS-E-Sim | **0.07** | **0.05** | **0.01** | **0.03** | **0.07** | **0.41** | **0.19** | 0.16 |
| WGAN | 0.08 | 0.04 | 0.02 | **0.03** | 0.06 | 0.39 | 0.21 | **0.15** |
| Original read (D-N) | 0.05 | 0.06 | 0.08 | 0.15 | 0.32 | 0.09 | 0.14 | 0.08 |
| DDS-E-Sim | **0.04** | **0.06** | 0.11 | 0.07 | 0.31 | 0.15 | 0.17 | 0.09 |
| WGAN | **0.04** | 0.07 | **0.09** | **0.09** | **0.32** | **0.14** | **0.16** | **0.08** |

#### Positional Skewness and Adaptive Stochasticity

To evaluate the possibility of overfitting or data leakage, we examine the positional error distributions as shown in Figures 2 and 3. These figures present both the total error counts and the base-specific error counts across sequence positions for insertions, deletions, and substitutions. The alignment of trends between the original datasets and the generated outputs demonstrates that our model avoids overfitting and data leakage. For dataset D-I, the generated sequences exhibit stochastic variation while maintaining overall consistency with the empirical error profiles (Figure 2). Specifically, insertion errors display terminal spikes, substitution errors are generally high, and deletion errors remain generally low except at terminal regions. Overall, the generated sequences successfully capture the error patterns observed in the original dataset.

**Figure 2.**
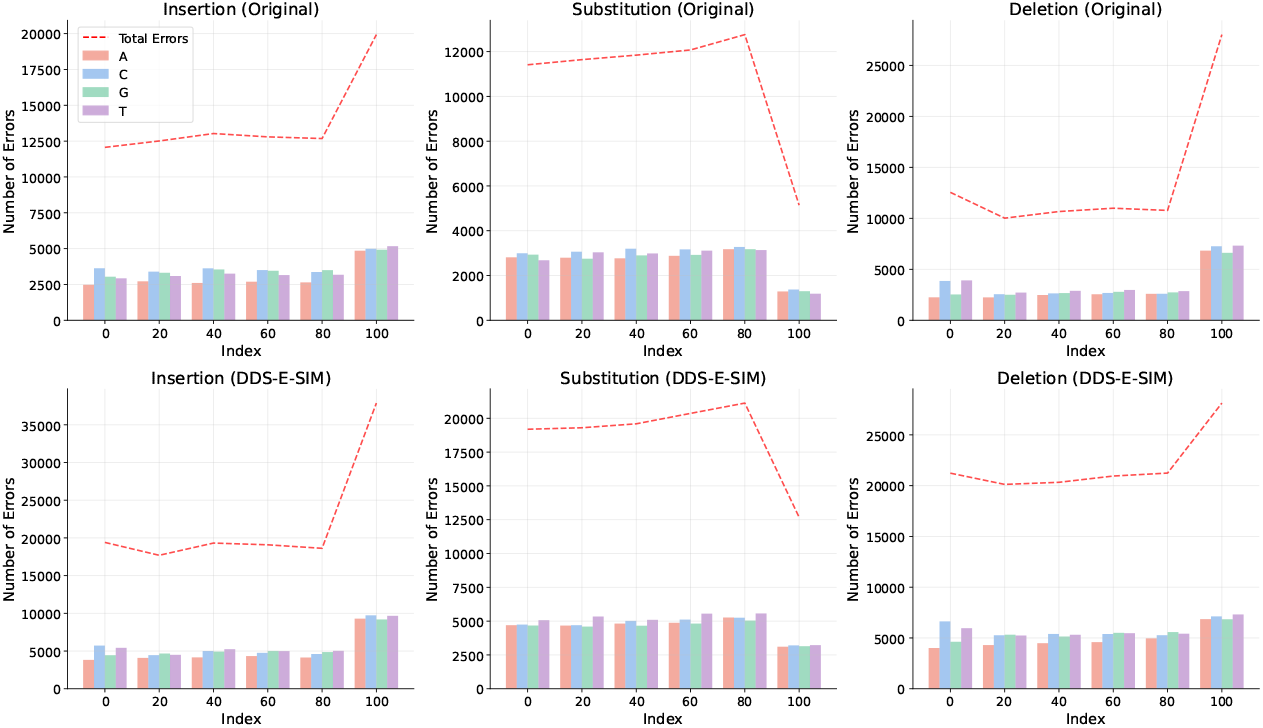
Stochastic insertion, substitution, and deletion error count in original and generated reads in dataset D-I. The x-axis denotes sequence position indices and the y-axis denotes error count. The bar plots represent base-specific error counts and the dotted line indicates the aggregate error count across all bases.

**Figure 3.**
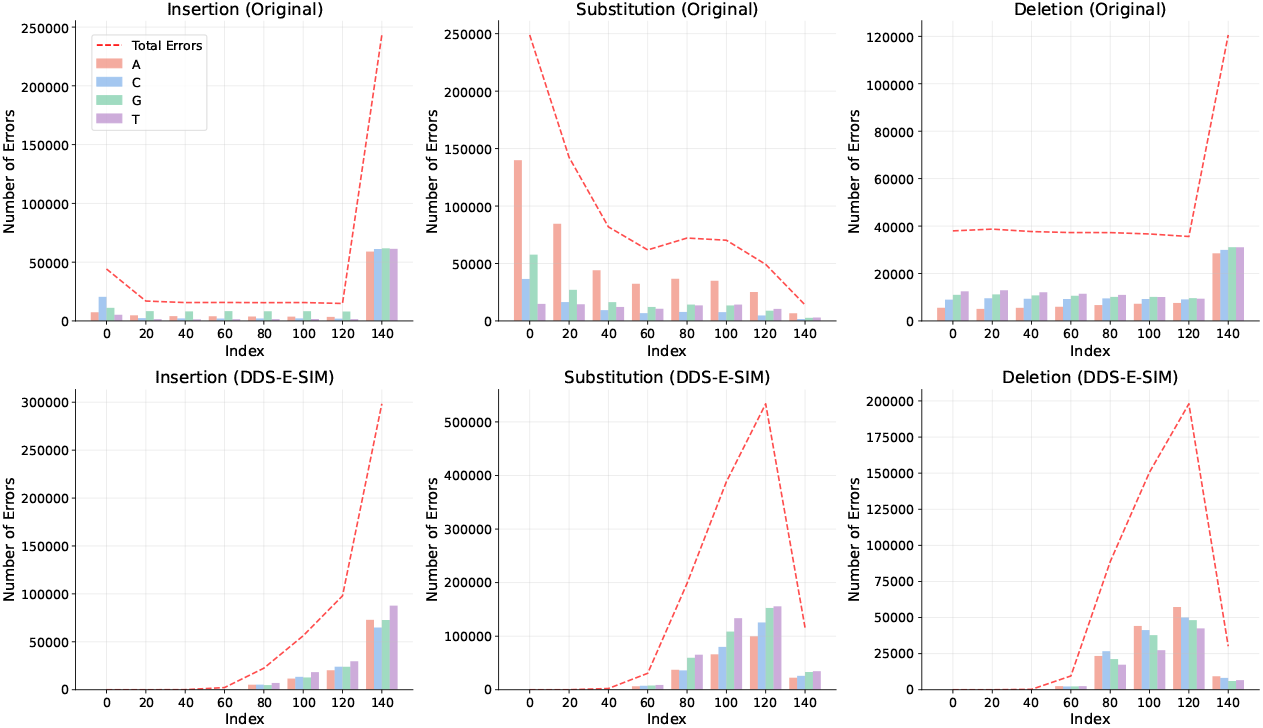
Stochastic insertion, substitution, and deletion error count in original and generated reads in dataset D-N. The x-axis denotes sequence position indices and the y-axis denotes error count. The bar plots represent base-specific error counts and the dotted line indicates the aggregate error count across all bases.

Similarly, we analyze positional distribution in the context of the D-N dataset as shown in Figure 3. Again, our model fairly matches the positional error distribution of the dataset except the distribution of substituition errors. In the D-N dataset, the substitution error rate is comparatively low, which is a probable reason for the noticeable mismatch in the overall error distribution.

#### Noise Injection for stochasticity

We experiment with adding noise in two ways to increase randomness in generated outputs at different coverages and prevent overfitting. In the first method, we add noise at the preprocessing stage of DNA sequences before they enter the local encoder. We observe an error rate of only 1.4%. In the second method, we introduce noise directly in the latent space of a Beta-VAE. We sample the latent variable *z* from a Gaussian distribution and add 3-5% noise to regularise the model. However, this method leads to an unexpectedly high error rate of 15.1%. The noise disrupts the latent space and prevents accurate sequence reconstruction. Thus, we proceed with the first method.

#### Stochasticity in DNA sequence reads

We demonstrated that DDS-E-Sim effectively captures the error patterns across multiple sequencing technologies. However, to faithfully mimic the DNA data storage pipeline, its output must also be stochastic; that is, it should generate diverse erroneous sequences for the same input while still following the underlying error distribution. To achieve this, we integrate a Beta-VAE into our architecture for stochastic read generation and further introduce Gaussian noise to the input sequences, which proves to be highly effective. To validate randomness, we generated reads from 35,329 fixed sequences at a coverage of 5 (simulating five reads per sequence), resulting in 100,743 unique oligos. This demonstrates that our model preserves not only the error distributions but also the essential stochasticity of the DNA data storage process.

## 4 Conclusion and Future Work

We present a universal transformer-based generative framework for simulating error-prone DNA sequences regardless of process or technology. The model captures higher-order error patterns while preserving stochasticity, achieving state-of-the-art performance. We are actively working to extend the framework with designing and combining stage-specific modules to simulate errors at each DDS stage and integrate error-correction algorithms for testing and optimization in the near future.

## A Appendix

### A.1 Additional Models Implementation

#### Multi-Attention-LSTM-Beta VAE (VAE)

We design a Beta-VAE architecture with Attention-LSTM for our ablation study to highlight the benefit of the transformer architectue in our propsed model. This architecture designed to efficiently handle high-dimensional data as shown in Figure 4. In the encoder, the DNA sequence is processed using Attention-LSTM, which effectively captures the complex error statistics in the DNA sequence. In the decoder, Attention-LSTM layers are used to model dependencies between positions in the DNA sequence. It preserves the sequence dynamics in the reconstructed error-prone output.

**Figure 4.**
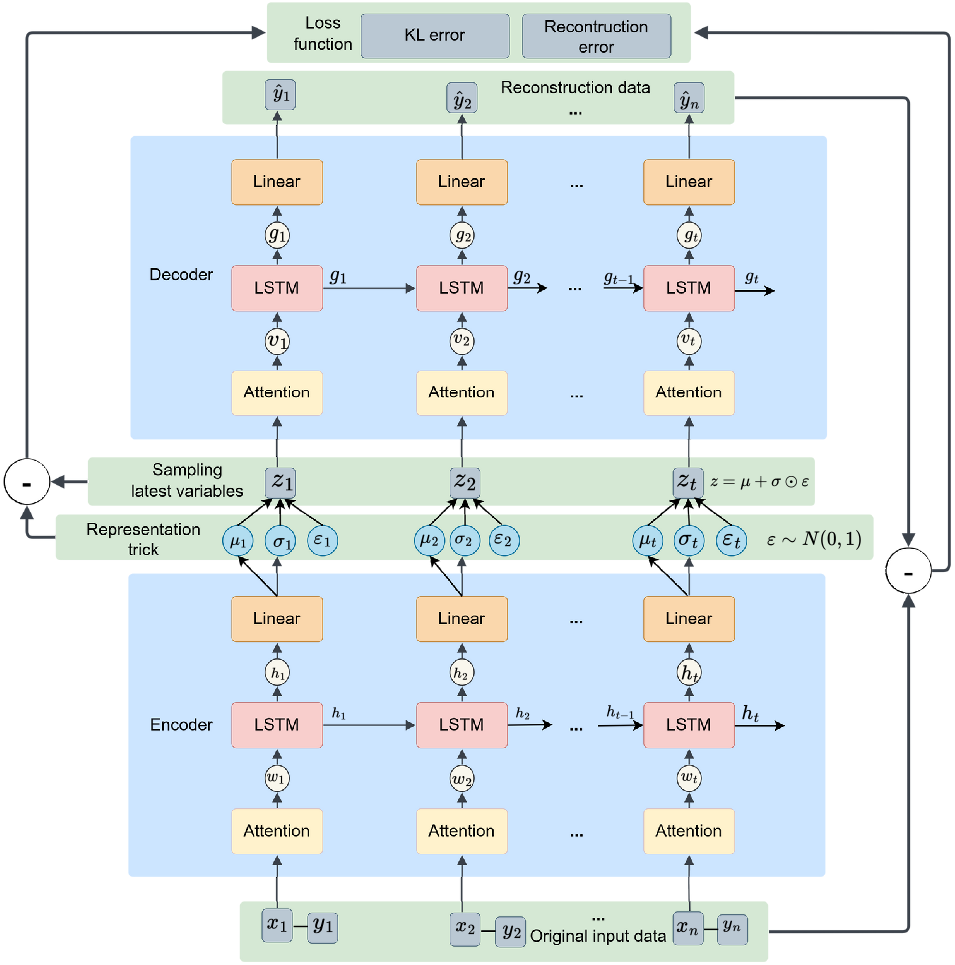
The architecture of the Attention-LSTM-VAE model

The input data is represented as *x* = [*x*_1_, *x*_2_, …, *x*_*n*_], where *x*_*n*_ denotes the nucleotide at the *n*-th position in the DNA sequence. The corresponding output data is represented as *y* = [*y*_1_, *y*_2_, …, *y*_*n*_], and the reconstructed error-prone output after the model processes the data is denoted as *ŷ* = [*ŷ*_1_, *ŷ*_2_, …, *ŷ*_*n*_]. During the encoding process, we obtain the mean *µ* = [*µ*_1_, *µ*_2_, …, *µ*_*n*_] and the variance *σ* = [*σ*_1_, *σ*_2_, …, *σ*_*n*_] that characterize the latent variables. The latent variables *z* = [*z*_1_, *z*_2_, …, *z*_*n*_] are then sampled, with *ϵ* = [*ϵ*_1_, *ϵ*_2_, …, *ϵ*_*n*_] representing the noise used in the reparameterization trick, where *ϵ* ∼ *N* (0, 1).

The key difference in Beta-VAE is the introduction of a beta parameter, *β*, that controls the trade-off between the reconstruction loss and the KL divergence. This parameter is introduced in the loss function to enforce a more structured and disentangled representation of the latent space. Thus, the KL divergence in the Beta-VAE is modified to:

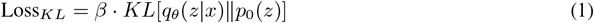

where *β* is a hyperparameter that determines the relative importance of the KL term compared to the reconstruction loss. When *β* = 1, the Beta-VAE becomes equivalent to the standard VAE. Increasing *β* enforces more disentanglement at the cost of reconstruction accuracy, while decreasing *β* gives more importance to reconstruction.

To obtain the posterior distribution of the latent variables *p*(*z*|*x*), we use variational inference. The true posterior distribution is approximated by a variational distribution *q*_*θ*_(*z*|*x*), which is parameterized by *θ*. The goal is to minimize the Kullback-Leibler (KL) divergence between the true posterior *p*(*z*|*x*) and the variational distribution *q*_*θ*_(*z*|*x*), which can be expressed as:

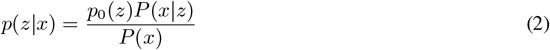

The KL divergence is computed as:

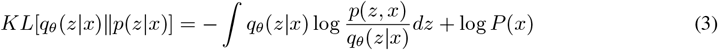

The Evidence Lower Bound (ELBO) is formulated as:

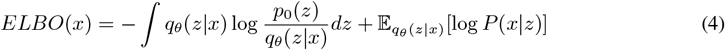

Minimizing the KL divergence is equivalent to maximizing the ELBO. We apply the reparameterization trick to allow gradient-based optimization. For the variational distribution *q*_*θ*_(*z* | *x*), we assume a Gaussian distribution with diagonal covariance:

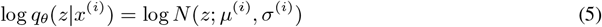

where *µ*^(*i*)^ and *σ*^(*i*)^ represent the mean and standard deviation for each input *x*^(*i*)^. Next, we compute the KL divergence between *q*_*θ*_(*z*|*x*) and the prior *N* (0, *I*):

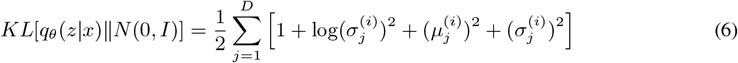

The objective function to optimize combines the modified KL divergence and the reconstruction error:

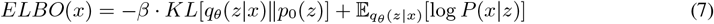

where the latent variables are sampled as:

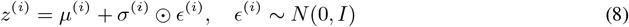

Here, ⊙ denotes element-wise multiplication between *σ*^(*i*)^ and *ϵ*^(*i*)^, and *ϵ*^(*i*)^ is sampled from a standard normal distribution.

The loss function for the Beta-VAE with Attention-LSTM is the sum of the KL divergence term and the reconstruction error, expressed as:

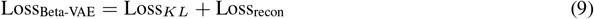

where the reparameterization is given by:

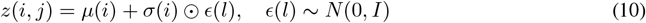

Beta-VAE with Attention-LSTM simplifies calculations despite its complexity. It boosts efficiency and robustness for handling noisy DNA storage data and error-prone sequence patterns.

#### Autoregressive Transformer (Transformer)

We utilize the Transformer architecture as shown in our proposed architecture in Figure 1 excluding the Beta-VAE framework for our ablation study.

### A.2 Additonal Implementation Details

#### DDS-E-Sim

We implement our model using PyTorch on an NVIDIA A100 GPU. We apply a padding of 5 to standardize input size. Using random search, we determine an optimal noise factor of 0.03 to induce stochasticity effectively. The model features an encoder with a 7-dimensional input, 256 model dimensions, 8 attention heads, 3 layers, a 128-dimensional latent space, a 1024 feedforward dimension, and a 0.1 dropout rate. The decoder mirrors this structure with a 7-dimensional output. We train our model with a batch size of 256 for 20 epochs using the Adam optimizer (learning rate = 1 × 10^−4^, *β* = 0.5).

#### VAE

We set the LSTM hidden size to 256 to capture complex dependencies. The output size is also 7, ensuring the model reconstructs the input. The sequence length is 154, representing the length of the input sequence. We handle the latent space with a latent dimension of 128. We divide the sequences into a few parts (usually 5) and feed that to the LSTMs as LSTMs cannot handle long sequences. We implement three LSTM layers in both the encoder and decoder. We train the model for 15 epochs.

